# Effectiveness of Digital Dispersion Compensation in OCT

**DOI:** 10.1101/2025.02.27.640677

**Authors:** Fengquan Liu, Xingde Li

## Abstract

Dispersion mismatch in optical coherence tomography (OCT) is typically addressed through either physical or digital compensation. In this study, we investigate the impact of dispersion on OCT detection sensitivity and compare the effectiveness of physical and digital compensation across varying degrees of dispersion mismatch. Our results demonstrate that while digital dispersion compensation can effectively restore detection sensitivity, its efficacy is constrained by the severity of the dispersion mismatch. Beyond a certain threshold, digital compensation fails to fully recover image information, leading to degradation in image quality.

## 1. Introduction

OCT has been established as a robust, noninvasive, and extremely valuable technology to visualize fine structures in biological tissues [1]. Dispersion mismatch between the reference and sample arms, however, induces point spread function (PSF) broadening and detection sensitivity (DS) drop, which compromises image resolution and contrast. There are in general two approaches to restore the PSF and the DS, known as physical dispersion compensation and digital dispersion compensation. The former commonly utilize gratings, prisms or GRISMs (gratings and prisms) [2] to anti-chirp the light, while the latter relies on post-experiment digital processing to sharpen the PSF [3-7]. However, it is unclear to what degree, digital dispersion remains effective. This question is particularly relevant when using a broadband, low-coherence light source with a central wavelength around 800 nm or within the visible range, as it is essential to determine the degree of fiber-optic length mismatch between the OCT reference and sample arms that can be tolerated [8-10]. In standard practice, optical fibers in both arms must be precisely cut to achieve closely matched fiber lengths, often requiring a tedious process, with any residual mismatch typically corrected using a pair of prisms inserted at an appropriate position [11, 12]. This paper presents a straightforward study comparing the DS of physically compensated and digitally compensated systems. The second section focuses on the theory of the effects of dispersion on the DS, while the comparison results are presented in the third section, demonstrating the effectiveness of digital compensation that affords to have optical fiber-length mismatch up to several centimeters.

## 2. Theoretical Analysis of the Impact of Dispersion Mismatch on Detection Sensitivity and OCT Axial Resolution

We first theoretically investigate the impact of dispersion mismatch on DS and PSF by comparing |*A*(*z*)|, the absolute value of the Fourier transforms of the collected interference signal *I*(*k*). For the case of DS comparison, since the systematic noise is consistent if the powers from both arms remain unchanged, the DS is proportional to the logarithm of the peak value of |*A*(*z*)|.

Consider a spectral-domain OCT system with a Gaussian source spectrum

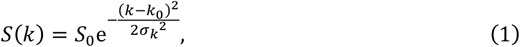

where *k*_0_ and *k* denotes the magnitude of center wave vector and wave vector in vacuum, respectively. Should a mirror be inserted in the sample arm, with the one-way optical pathlength difference between the sample and reference arm denoted as *z*_0_, the collected interference signal can be written as

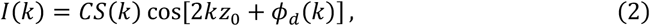

where *ϕ*_*d*_(*k*) represents the high-order dispersive phase induced by dispersion mismatch between the two arms, and *C* is a dispersion-independent constant that is trivial in the following discussion and will thus be dropped. For simplicity, we consider up to second-order dispersion, and the dispersive phase should thus take the form:

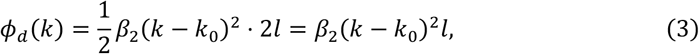

where *l* is the one-way optical material mismatch between the sample and reference arm, and *β*_2_ is the second-order dispersion coefficient of the material. For compactness only one dispersive material is included, but it will be shown that the conclusion of this section can be generalized to the case where more than one dispersive material is involved.

The Fourier transform of the interference signal (Eq. (2)) can now be expressed as

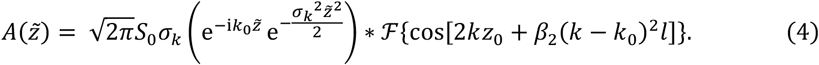

where ∗ denotes the convolution operation, 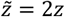 where *z* is the relative optical pathlength. We are only interested in |*A*(*z* ≥ 0)|, and given that for general cases 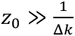

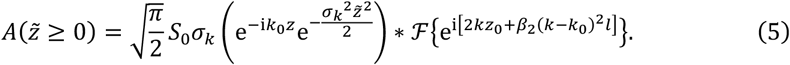

Subsequently (deduction provided in Supplemental Document), performing the convolution, we arrive at

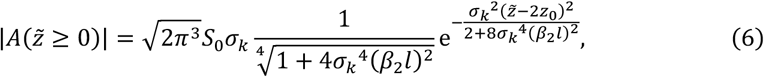

from which we obtain the peak value:

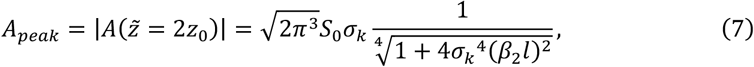

and therefore, the DS difference between a dispersive system and a dispersion-free (or dispersion matched) system could be written as

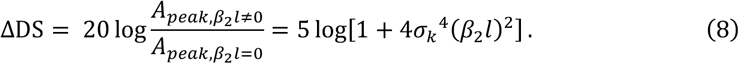

As throughout the deduction process, the product *β*_2_*l* has always been treated holistically, *β*_2_*l* in the equations above can be generalized and replaced by ∑_*i*_ *β*_2,*i*_*l*_*i*_ , in which the subscript *i* denotes the type of dispersive materials in the system. We would also be able to obtain the FWHM of the PSF from Eq. (7):

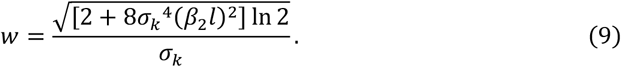

Note that the value *w* in Eq. (9) is half of the FWHM of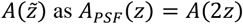.

To validate the theoretical model, we performed experiments using a spectral domain OCT system [13]. Mirrors were placed in both the reference and sample arms with material balanced neutral density (ND) filters placed in both arms to avoid detector saturation. A pair of BK7 prisms was inserted in the reference arm for adjusting dispersion mismatch. The system was first optimized by fine-tuning the insertion length of BK7 to minimize the FWHM of PSF, in which case the system was considered physically compensated. The length of BK7 in the optical path was then altered to add dispersion to the system, in the process of which the DS and FWHM of PSF were calculated. These measured values were compared with the theoretical values, where *σ*_*k*_ was related to the Gaussian spectral bandwidth as defined in Eq. (1). As the actual spectrum shape of the light source deviates from perfect Gaussian, small discrepancies between theoretical and measured values were observed, as demonstrated in Figure 1. The general dependence, however, is in excellent agreement, with *R*^2^ = 0.95 for the detection sensitivity model and *R*^2^ = 0.99 for the PSF model.

**Fig. 1.**
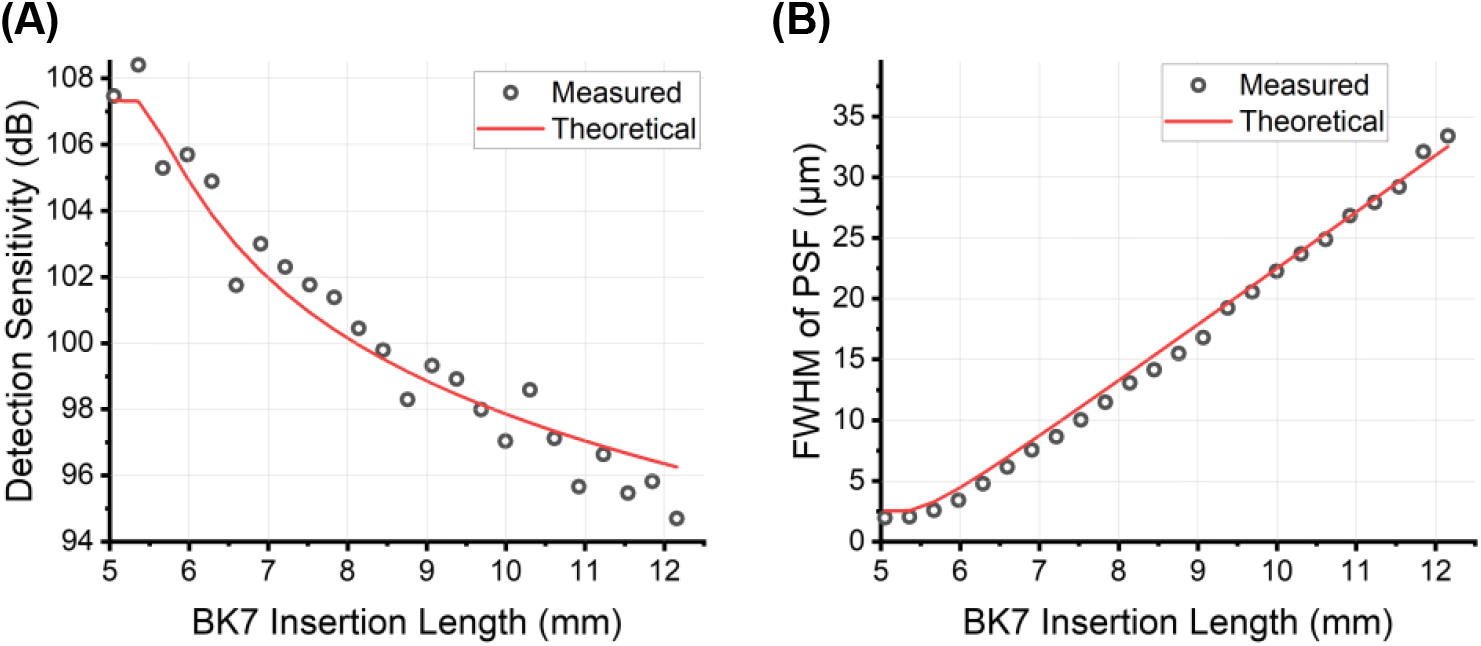
Impact of dispersion on (A) detection sensitivity and (B) full width at half maximum (FWHM) of the point spread function (PSF).

## 3. Experimental Demonstration of the Effectiveness of Digital Dispersion Compensation

This section presents the DS comparison between a dispersive system, a digitally compensated system, and a physically compensated system. A standard digital dispersion compensation algorithm [4, 14] is implemented in this study, with the basic scheme as follows. The OCT interference signal *I*(*k*) shown in Eq. (2) was first filtered and Hilbert transformed to obtain the phase *ϕ*(*k*). The high-order dispersive phase *ϕ*_*d*_(*k*) was then extracted, leaving the linear component of the phase and the trivial zeroth-order constant. The Hilbert-transformed signal was then multiplied by 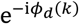 to compensate for dispersion mismatch and the outcome was then dispersion-compensated. The DS and the FWHM of the compensated PSF were then calculated, as shown respectively in Figure 2(A) and (B).

**Fig. 2.**
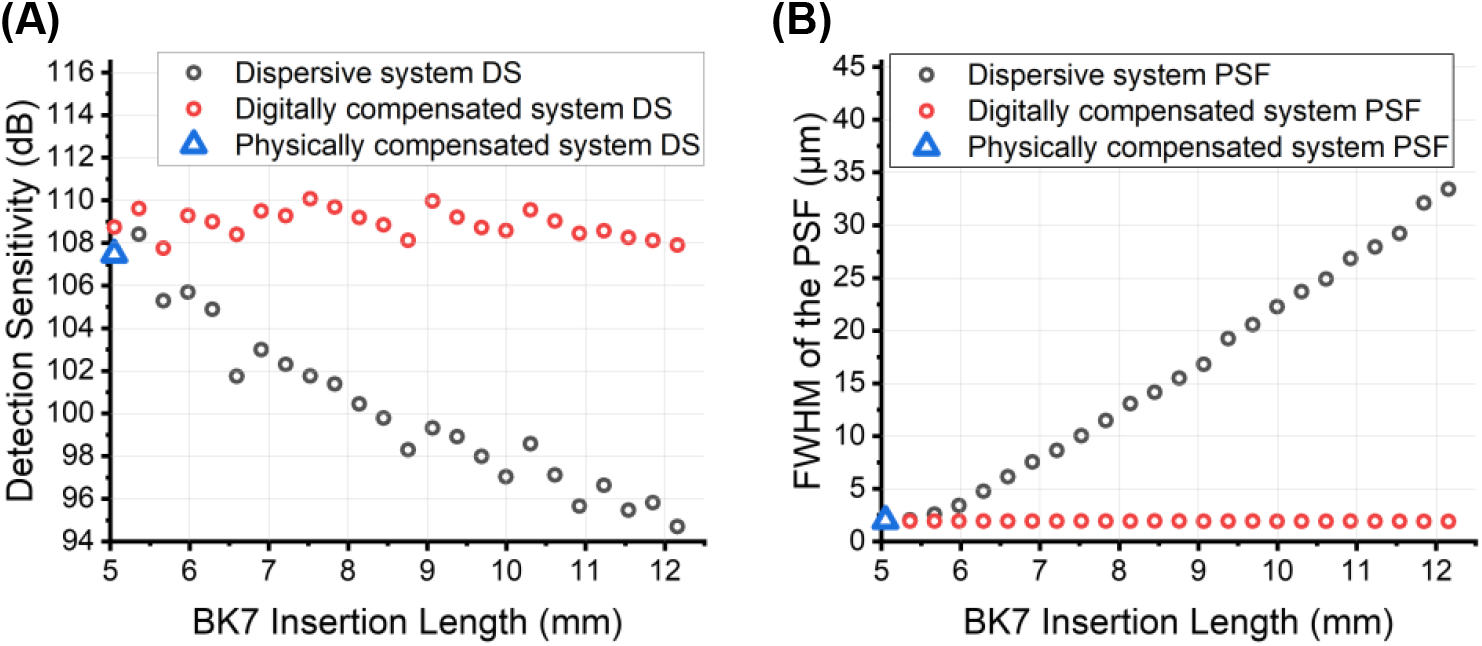
Comparison between (A) detection sensitivity (DS) and (B) full width at half maximum (FWHM) of the point spread function (PSF) for the dispersive system (black circles), the digitally compensated system (red circles) and the physically compensated system (blue triangle).

Digital dispersion compensation has demonstrated robust performance in highly dispersive systems by effectively restoring the DS (Figure 2(A)) and recovering the ideal PSF (Figure 2(B)). Digital compensation even further improved the DS over the physically compensated system by 1 dB. This improvement might arise from the inherent limitation of using only a single prism pair for physical compensation, which makes it challenging to fully compensate for all orders of dispersion. Digital compensation, on the other hand, inherently removes all orders of dispersion, thereby further optimizing the DS beyond the capabilities of physical compensation.

Given the critical role of DS in image contrast, we compared images acquired using a physically compensated system, dispersive systems with fiber length mismatches of up to 100 mm, and the digitally compensated version of the dispersive system. The physically compensated system was tuned as described in Section 2, achieving a PSF FWHM of 2.3 µm. PSFs were collected prior to imaging for subsequent digital compensation. For *in vivo* imaging, a mouse was first anesthetized using 4.5% isoflurane with supplemental oxygen and underwent hair removal procedures. The mouse was then maintained under anesthesia with 1.5% isoflurane and supplemental oxygen while a 2 mm × 2 mm region of the dorsal skin was imaged. Animal experimental procedures were approved by the Animal Care and Use Committee at the Johns Hopkins University. Our findings indicate that digital dispersion compensation successfully restored DS to the level of the physically compensated system for fiber length mismatches of up to 40 mm, corresponding to a severely deteriorated PSF FWHM of 183 µm, as shown by the red curve in Figure 3(A). The digitally compensated images closely resemble those acquired with the physically compensated system, exhibiting no noticeable loss in contrast, imaging depth, or resolution, as demonstrated in Figures 3(B-D). Notably, while the dispersive image appears blurred, lacking clear boundaries and discernible tissue layer details, digital dispersion compensation effectively restores tissue fine structures. The digitally compensated image clearly delineates the epidermis, the DEJ (dermal-epidermal junction), the boundaries of adipose tissue and underlying muscle, as well as the individual adipose droplets. The minor discrepancies between Figures 3(B) and 3(C) may be attributed to motion artifacts from the mouse during *in vivo* imaging.

**Fig. 3.**
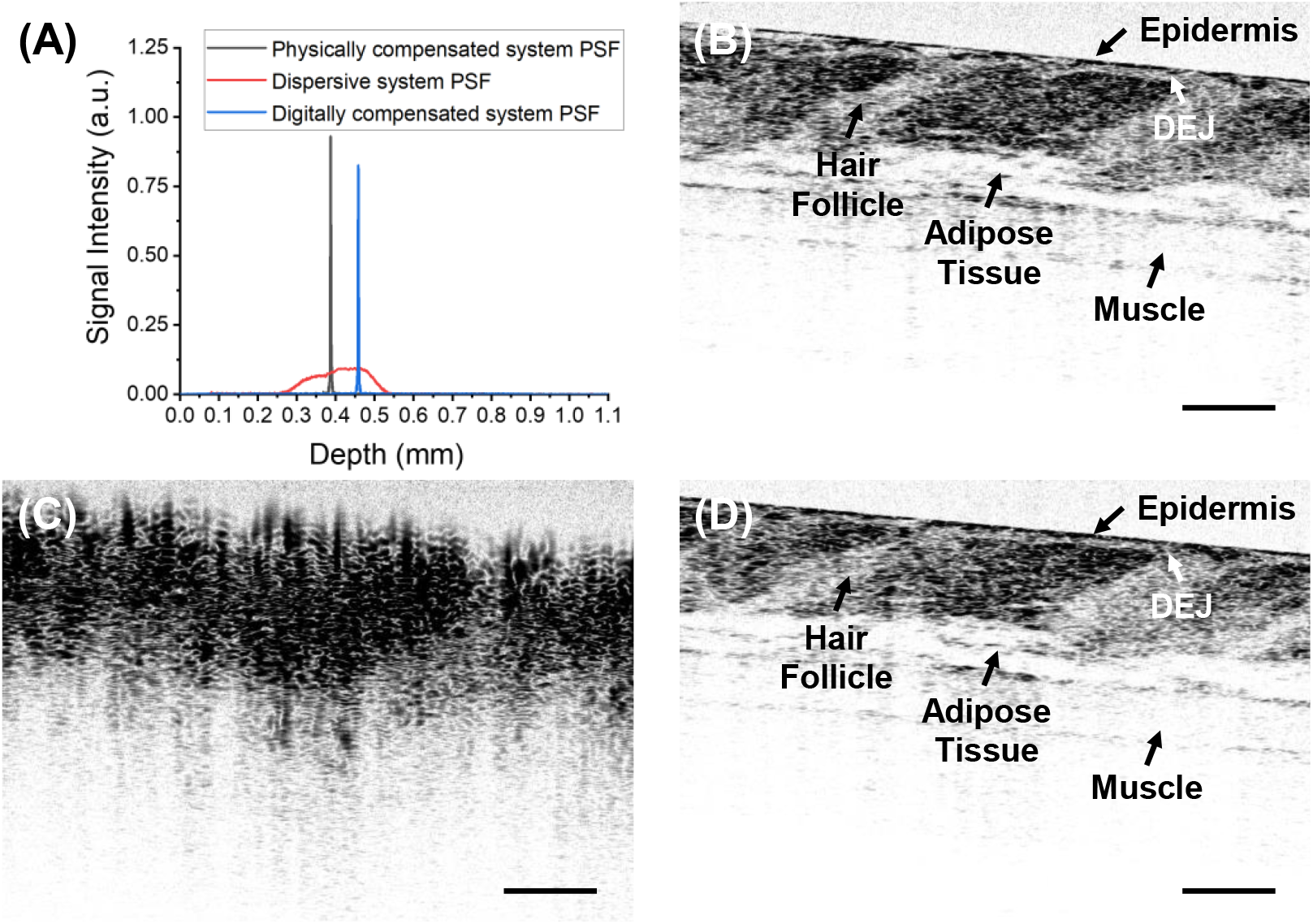
(A) PSFs of the physically compensated system (black), the dispersive system (red) and the digitally compensated system (blue), and *in vivo* mouse skin tissue image taken with (B) the physically compensated system, (C) the dispersive system with 40 mm fiber length mismatch, and (D) the digitally compensated system. DEJ: dermal-epidermal junction. Scale bars: 100 µm.

However, at a fiber length mismatch of 100 mm, corresponding to a very poor PSF FWHM of 580 µm as shown in Figure 4(A), digital dispersion compensation is no longer able to fully restore DS. A 7 dB reduction in DS, compared to the physically compensated system, was observed in the digitally compensated system. As a result, the digitally compensated image exhibits a decline in contrast and a loss of deep-tissue information. As is demonstrated in Figures 4(B-D), the muscle, while being clearly visible in the image taken with the physically compensated system, is unrecognizable in the digitally compensated image. These findings suggest that digital dispersion compensation has inherent limitations, and excessive dispersion beyond a certain threshold leads to irreversible loss of image information, as deep-tissue signal becomes so dispersed that it is buried beneath the noise level and unrecoverable. It is important to note that this limitation can be case-dependent, as several factors, including the power of the OCT light source, system roll-off, optical characteristics of the sample arm probe, the properties of the tissue itself, and ultimately the overall system noise level, collectively influence the robustness of digital compensation against excessive dispersion.

**Fig. 4.**
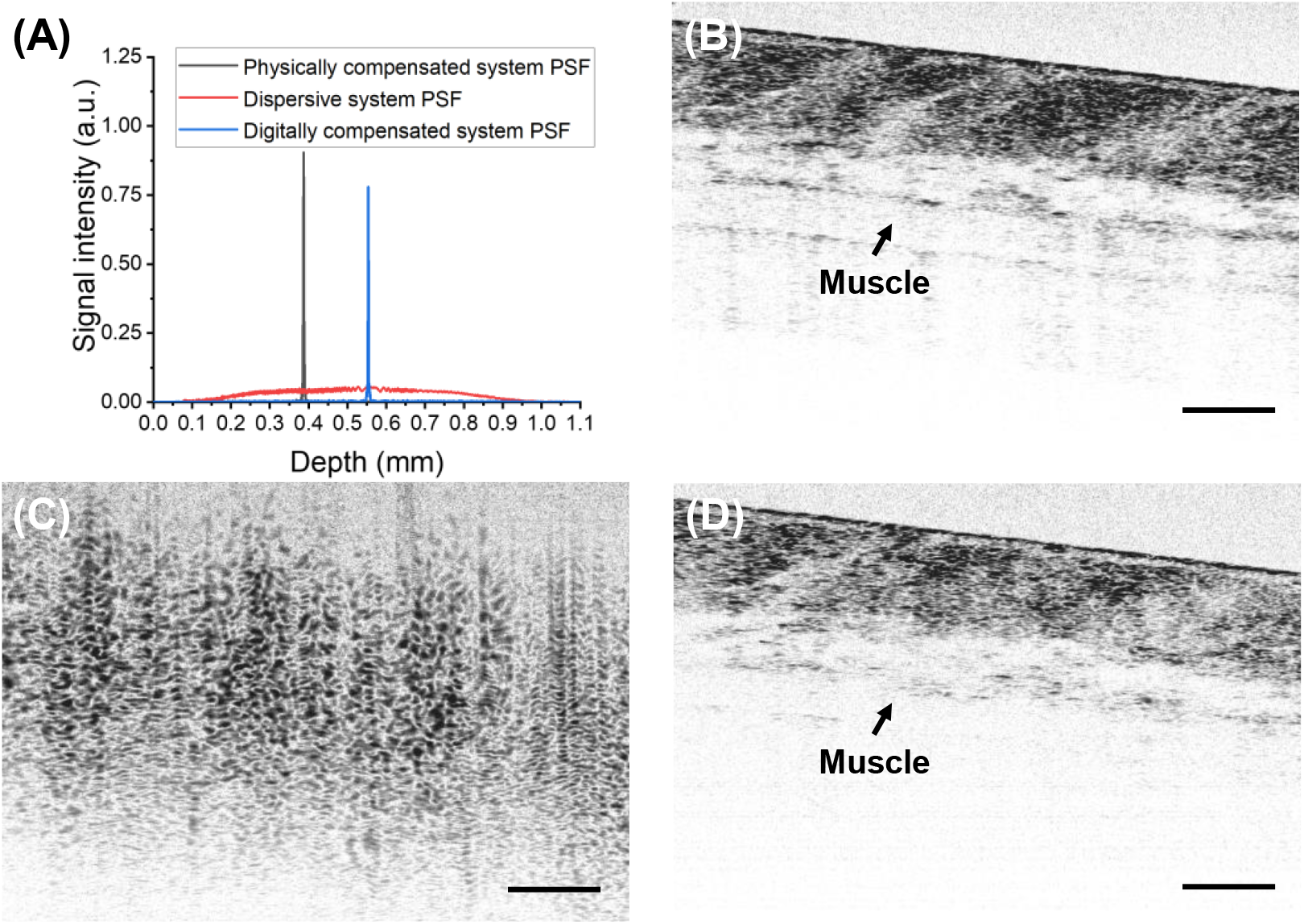
(A) PSFs of the physically compensated system (black), the dispersive system (red) and the digitally compensated system (blue), and *in vivo* mouse skin tissue image taken with (B) the physically compensated system, (C) the dispersive system with 100 mm fiber length mismatch, (D) the digitally compensated system. Scale bars: 100 µm.s

## 4. Conclusion

We have developed a theoretical model describing the impact of dispersion on detection sensitivity and compared the PSFs and images obtained from dispersive, digitally compensated, and physically compensated systems. The results demonstrate that while dispersion mismatch in OCT significantly degrades the detection sensitivity of the system, digital dispersion compensation can afford a fiber-length mismatch up to 4 cm (or even more) between the OCT reference and sample arms and can effectively restore detection sensitivity to levels comparable to those achieved with physical compensation. However, when dispersion mismatch exceeds a certain range, deep-tissue image information is lost, and the compromised image contrast can no longer be effectively rescued by digital compensation.

## Funding

National Institutes of Health (NIH) (R01 EB033364, P41 EB032840).

## Acknowledgment

The authors thank Cheng-Yu Lee for assistance with data collection.

## Notes

### Competing Interest Statement

The authors have declared no competing interest.

## References

1. D. Huang, E.A. Swanson, C.P. Lin, et al., “Optical Coherence Tomography.” Science 254(5035), 1178–1181 (1991).

2. W. Liang, G. Hall, and X. Li, “Spectro-temporal dispersion management of femtosecond pulses for fiber-optic two-photon endomicroscopy.” Optics Express 26(18), 22877–22893 (2018).

3. D.L. Marks, A.L. Oldenburg, J.J. Reynolds, et al., “Digital algorithm for dispersion correction in optical coherence tomography for homogeneous and stratified media.” Applied Optics 42(2), 204–217 (2003).

4. B. Cense, N.A. Nassif, T.C. Chen, et al., “Ultrahigh-resolution high-speed retinal imaging using spectral-domain optical coherence tomography.” Optics Express 12(11), 2435–2447 (2004).

5. M. Wojtkowski, V.J. Srinivasan, T.H. Ko, et al., “Ultrahigh-resolution, high-speed, Fourier domain optical coherence tomography and methods for dispersion compensation.” Optics Express 12(11), 2404–2422 (2004).

6. N. Lippok, S. Coen, P. Nielsen, et al., “Dispersion compensation in Fourier domain optical coherence tomography using the fractional Fourier transform.” Optics Express 20(21), 23398–23413 (2012).

7. A. Kho, and V.J. Srinivasan, “Compensating spatially dependent dispersion in visible light OCT.” Optics Letters 44(4), 775–778 (2019).

8. W. Drexler, U. Morgner, F.X. Kärtner, et al., “In vivo ultrahigh-resolution optical coherence tomography.” Optics Letters 24(17), 1221–1223 (1999).

9. K. Li, W. Liang, J. Mavadia-Shukla, et al., “Super-achromatic optical coherence tomography capsule for ultrahigh-resolution imaging of esophagus.” Journal of Biophotonics 12(3), e201800205 (2019).

10. J. Yi, Q. Wei, W. Liu, et al., “Visible-light optical coherence tomography for retinal oximetry.” Optics Letters 38(11), 1796–1798 (2013).

11. J. Mavadia-Shukla, P. Fathi, W. Liang, et al., “High-speed, ultrahigh-resolution distal scanning OCT endoscopy at 800 nm for in vivo imaging of colon tumorigenesis on murine models.” Biomedical Optics Express 9(8), 3731–3739 (2018).

12. H.-C. Park, D. Li, R. Liang, et al., “Multifunctional Ablative Gastrointestinal Imaging Capsule (MAGIC) for Esophagus Surveillance and Interventions.” BME Frontiers 5, 0041 (2024).

13. W. Yuan, J. Mavadia-Shukla, J. Xi, et al., “Optimal operational conditions for supercontinuum-based ultrahigh-resolution endoscopic OCT imaging.” Optics Letters 41(2), 250–253 (2016).

14. J. Wang, C. Xu, S. Zhu, et al., “A Generic and Effective System Dispersion Compensation Method: Development and Validation in Visible-Light OCT.” Photonics 10(8), 892 (2023).

