## Supplemental Document for "Effectiveness of Digital Dispersion Compensation in OCT"

### Effectiveness of Digital Dispersion Compensation in OCT: Supplemental Document

##### Deduction from Eq. (5) to Eq. (6).

Eq. (5) is given as:

$$A(\tilde{z} \geq 0) = \sqrt{\frac{\pi}{2}} S_0 \sigma_k \left( \mathrm{e}^{-\mathrm{i}k_0\tilde{z}} \mathrm{e}^{-\frac{\sigma_k{}^2 \tilde{z}^2}{2}} \right) * \mathcal{F}\left\{ \mathrm{e}^{\mathrm{i}\left[2kz_0 + \beta_2 (k-k_0)^2 l\right]} \right\}, \tag{S1}$$

where

$$\begin{aligned}
\mathcal{F}\left\{ \mathrm{e}^{\mathrm{i}\left[2kz_0 + \beta_2 (k-k_0)^2 l\right]} \right\} &= \int_{-\infty}^{+\infty} \mathrm{e}^{\mathrm{i}\left[2kz_0 + \beta_2 (k-k_0)^2 l\right]} \mathrm{e}^{-\mathrm{i}k\tilde{z}} \, \mathrm{d}k \\
&= \mathrm{e}^{\mathrm{i}k_0(2z_0 - \tilde{z})} \int_{-\infty}^{+\infty} \mathrm{e}^{\mathrm{i}\left[2(k-k_0)z_0 + \beta_2 (k-k_0)^2 l\right]} \mathrm{e}^{-\mathrm{i}(k-k_0)\tilde{z}} \, \mathrm{d}k \\
&= \mathrm{e}^{\mathrm{i}k_0(2z_0 - \tilde{z})} \int_{-\infty}^{+\infty} \mathrm{e}^{\mathrm{i}\left[\beta_2 l \kappa^2 + (2z_0 - \tilde{z})\kappa\right]} \, \mathrm{d}\kappa \\
&= \mathrm{e}^{\mathrm{i}k_0(2z_0 - \tilde{z})} \mathrm{e}^{-\mathrm{i}\frac{(2z_0 - \tilde{z})^2}{4\beta_2 l}} \int_{-\infty}^{+\infty} \mathrm{e}^{\mathrm{i}\beta_2 l \left[\kappa + \frac{(2z_0 - \tilde{z})}{2\beta_2 l}\right]^2} \, \mathrm{d}\kappa \\
&= \mathrm{e}^{\mathrm{i}k_0(2z_0 - \tilde{z})} \mathrm{e}^{-\mathrm{i}\frac{(2z_0 - \tilde{z})^2}{4\beta_2 l}} \cdot \sqrt{\frac{\pi}{2\beta_2 l}} (1 + \mathrm{i}) \,,
\end{aligned} \tag{S2}$$

in which

$$\int_{-\infty}^{+\infty} \mathrm{e}^{\mathrm{i}x^2} \, \mathrm{d}x = \sqrt{\frac{\pi}{2}} (1 + \mathrm{i}) \tag{S3}$$

is utilized at the last equal sign.

Substituting the expression in Eq. (S2) to Eq. (S1) yields:

$$A(\tilde{z} \geq 0) = (1 + \mathrm{i}) \sqrt{\frac{\pi^2}{4\beta_2 l}} S_0 \sigma_k \left[ \left( \mathrm{e}^{-\mathrm{i}k_0\tilde{z}} \mathrm{e}^{-\frac{\sigma_k{}^2 \tilde{z}^2}{2}} \right) * \left( \mathrm{e}^{\mathrm{i}k_0(2z_0 - \tilde{z})} \mathrm{e}^{-\mathrm{i}\frac{(2z_0 - \tilde{z})^2}{4\beta_2 l}} \right) \right], \tag{S4}$$

where

$$\begin{aligned}
|F(\tilde{z})| &\equiv \left| \left( \mathrm{e}^{-\mathrm{i}k_0\tilde{z}} \mathrm{e}^{-\frac{\sigma_k{}^2 \tilde{z}^2}{2}} \right) * \left( \mathrm{e}^{\mathrm{i}k_0(2z_0 - \tilde{z})} \mathrm{e}^{-\mathrm{i}\frac{(2z_0 - \tilde{z})^2}{4\beta_2 l}} \right) \right| \\
&= \left| \int_{-\infty}^{+\infty} \mathrm{e}^{-\mathrm{i}k_0\zeta} \mathrm{e}^{-\frac{\sigma_k{}^2 \zeta^2}{2}} \mathrm{e}^{-\mathrm{i}k_0(\tilde{z} - 2z_0 - \zeta)} \mathrm{e}^{-\mathrm{i}\frac{(\tilde{z} - 2z_0 - \zeta)^2}{4\beta_2 l}} \, \mathrm{d}\zeta \right| \\
&= \left| \mathrm{e}^{-\mathrm{i}k_0(\tilde{z} - 2z_0)} \right| \left| \int_{-\infty}^{+\infty} \mathrm{e}^{-\frac{\sigma_k{}^2 \zeta^2}{2}} \mathrm{e}^{-\mathrm{i}\frac{(\tilde{z} - 2z_0 - \zeta)^2}{4\beta_2 l}} \, \mathrm{d}\zeta \right|.
\end{aligned} \tag{S5}$$

For ease of discussion, we set $a = \frac{\sigma_k{}^2}{2}$, $b = \frac{1}{4\beta_2 l}$ and $z_p = \tilde{z} - 2z_0$. The second term on the side of the last equal sign of Eq. (S5) can now be written as

$$\left| \int_{-\infty}^{+\infty} \mathrm{e}^{-a\zeta^2 - \mathrm{i}b\left(z_p - \zeta\right)^2} \, \mathrm{d}\zeta \right| = \left| \int_{-\infty}^{+\infty} \mathrm{e}^{-(a + \mathrm{i}b)\zeta^2 + 2\mathrm{i}bz_p\zeta - \mathrm{i}bz_p^2} \, \mathrm{d}\zeta \right|$$

$$= \left| e^{-ibz_p^2} e^{-\frac{b^2 z_p^2}{a+ib}} \int_{-\infty}^{+\infty} e^{-(a+ib)\left(\zeta - \frac{ibz_p}{a+ib}\right)^2} d\zeta \right|$$
$$= e^{-\frac{ab^2 z_p^2}{a^2+b^2}} \left| \int_{-\infty + i\frac{abz_p}{a^2+b^2}}^{+\infty + i\frac{abz_p}{a^2+b^2}} e^{-(a+ib)\xi^2} d\xi \right|. \quad (S6)$$

where a substitution of $\xi = \zeta - \frac{ibz_p}{a+ib}$ is made. Utilizing a rectangular contour and the Cauchy integral theorem, as well as the fact that the integrand is symmetric with respect to the imaginary axis in the complex plane, we have

$$\left| \int_{-\infty + i\frac{abz_p}{a^2+b^2}}^{+\infty + i\frac{abz_p}{a^2+b^2}} e^{-(a+ib)\xi^2} d\xi \right| = \left| \int_{-\infty}^{+\infty} e^{-(a+ib)\xi^2} d\xi \right|. \quad (S7)$$

Set $a + ib = re^{i\theta}$, where $r = \sqrt{a^2 + b^2}, \theta = \tan^{-1}\frac{b}{a}$. The right-hand side of Eq. (S7) can then be expressed as

$$\left| \int_{-\infty}^{+\infty} e^{-(a+ib)\xi^2} d\xi \right| = \left| \int_{-\infty}^{+\infty} e^{-re^{i\theta}\xi^2} d\xi \right|. \quad (S8)$$

Consider a sector contour in the complex plane with central angle $\frac{\theta}{2}$ and radius $R \to +\infty$. The integral on the arc of the sector is zero is trivial and the Cauchy integral theorem yields:

$$\left| \int_{-\infty}^{+\infty} e^{-re^{i\theta}\xi^2} d\xi \right| = 2 \left| e^{-i\frac{\theta}{2}} \int_{0}^{+\infty} e^{-r\xi^2} d\xi \right| = \sqrt{\frac{\pi}{r}} = \frac{\sqrt{\pi}}{\sqrt[4]{a^2 + b^2}}. \quad (S9)$$

Plugging the equations above to Eq. (S6) yields

$$\left| \int_{-\infty}^{+\infty} e^{-a\zeta^2 - ib(z_p - \zeta)^2} d\zeta \right| = \frac{\sqrt{\pi}}{\sqrt[4]{a^2 + b^2}} e^{-\frac{ab^2 z_p^2}{a^2+b^2}} = \frac{\sqrt{\pi}}{\sqrt[4]{\frac{\sigma_k{}^4}{4} + \frac{1}{16(\beta_2 l)^2}}} e^{-\frac{\sigma_k{}^2(\tilde{z}-2z_0)^2}{2+8\sigma_k{}^4(\beta_2 l)^2}}. \quad (S10)$$

and further substitution to Eq. (S5) and (S4) yields

$$|A(\tilde{z} \geq 0)| = \sqrt{\frac{\pi^2}{2\beta_2 l}} S_0 \sigma_k |F(\tilde{z})| = \sqrt{2\pi^3} S_0 \sigma_k \frac{1}{\sqrt[4]{1 + 4\sigma_k{}^4(\beta_2 l)^2}} e^{-\frac{\sigma_k{}^2(\tilde{z}-2z_0)^2}{2+8\sigma_k{}^4(\beta_2 l)^2}}, \quad (S11)$$

which is exactly Eq. (6).
